# BayesR3AD: Joint analysis of additive and dominance in Bayesian mixture models

**DOI:** 10.64898/2026.02.23.707560

**Authors:** Huan Yuan, Edmond J. Breen, Iona M. MacLeod, Majid Khansefid, Ruidong Xiang, Michael E. Goddard

## Abstract

**Background:** Genomic prediction in livestock is predominantly based on additive models, even though dominance and other non-additive effects can contribute appreciably to phenotypic variance for fitness and fertility traits. Bayesian mixture models, such as Bayes R, have proven effective for modelling sparse, heterogeneous additive SNP effects, but most implementations do not explicitly accommodate dominance. In this study, we extended BayesR3 to jointly model additive and dominance marker effects within a unified Bayesian mixture framework, denoted BayesR3AD, and used this method to estimate additive and dominance effects for fertility and cow survival (longevity) in Holstein cattle.

**Results:** Using real Holstein genotypes (227,942 animals, 74,626 SNPs), we simulated phenotypes with additive and dominance effects. When dominance was present in the simulated data, BayesR3AD improved prediction accuracy of genetic values by +0.1011 (0.6144 vs 0.5133; ≈19.7% relative) compared with the additive-only BayesR3 model and recovered additive and dominance variance components without bias. Under purely additive simulations, dominance mixture components were effectively empty, confirming that the extended model shrinks unnecessary dominance effects toward zero. In real fertility data, including calving interval (63,378 records) and survival (68,514 records), BayesR3AD estimated small dominance variance (≈1-3% of total genetic variance). The model highlighted a very large additive loci at 57.82 Mb on BTA18 for both calving interval and survival. concordant with previous GWAS studies of Holstein fertility. Additionally, a large dominance effect was found at 44.37 Mb on BTA18 for calving interval implicating a heterozygote advantage that increases fertility.

**Conclusions:** BayesR3AD provides a practical extension of BayesR3 that captures both additive and dominance contributions to genomic prediction. The method is robust, reverting effectively to the additive model when dominance is absent, while delivering accurate variance decomposition, and potential gains in prediction accuracy when dominance is present. Application to Holstein fertility traits demonstrates that dominance can be detected and quantified without compromising additive inference, supporting improved prediction of total genetic merit. While validated in cattle, BayesR3AD can be directly applied to other species to better model and predict traits related to fitness.

## Background

The introduction of dense genome-wide SNP arrays has enabled genomic selection in livestock and crop improvement, leading to substantial gains in genetic progress and reductions in generation interval [1-3]. Dairy cattle have been the focus of extensive genomic evaluation efforts, with large reference populations and routinely recorded phenotypes for production, health, and fertility traits [1, 4]. Most genomic prediction pipelines in this context are based on additive linear mixed models such as Genomic Best Linear Unbiased Prediction (GBLUP), or on Bayesian regression models that assume marker effects follow a long tailed distribution often described by a mixture distribution [3, 5]. Bayesian mixture models [3] explicitly implement a long tailed distribution of effects by assigning each variant effect to one of several normal distributions with different variances, including a spike at zero [6]. This flexibility can improve both mapping and prediction performance by allowing the data to inform the sparsity and distribution of effect sizes [7, 8]. The BayesR3 implementation further introduces algorithmic refinements and computational improvements that enable efficient large-scale genomic prediction, SNP-effect inference, and fine mapping of putative causal variants [9].

Despite the widespread success of additive genomic prediction models, non-additive genetic effects are well known, particularly for traits related to fitness (e.g., [10-14]). Ignoring dominance can bias estimates of additive variance and misattribute non-additive signal to the residual, ultimately reducing the accuracy of total genetic merit predictions [15]. It may also obscure important biological mechanisms, such as recessive deleterious alleles and heterozygosity-related effects [16]. Although several authors have extended GBLUP frameworks to incorporate dominance relationship matrices [12, 13], there is comparatively less work on integrating dominance within Bayesian mixture models that estimate SNP effects directly, enabling sparse, locus-specific modelling and clearer biological interpretation [6, 7].

The goal of this paper is to extend BayesR3 to an integrated additive-dominance model. In this paper, the additive-only specification is denoted BayesR3, whereas the extended formulation that incorporates both additive and dominance effects is denoted BayesR3AD. We specify genotype coding for additive and dominance components, define mixture priors for SNP effects, derive the full conditional distributions used in the Gibbs sampler, and explain how to decompose genetic variance into additive and dominance parts. We show that BayesR3AD correctly estimates variant effects and variances using simulated data and then apply it to data on calving interval and survival in Holstein dairy cows.

## Methods

### Data

For the simulation study, we used real genotypes of 227,942 Australian Holstein dairy cows. The genotypes included 74,626 SNPs from a custom SNP panel used for genomic evaluations in Australia, which was described in van den Berg et al. (2024) [17]. The samples were randomly partitioned into a training set (70%, N=159,559) and a validation set (30%, N=68,383). Phenotypes were simulated to have the broad-sense heritability H^2^ ≈ 0.44. Details of the simulation model used to generate phenotypes for these cows (with and without dominance effects) are described in the later section of Methods.

For real-data analyses, we used Holstein cow data from DataGene’s April 2025 national evaluation, which had 63,378 records on a fertility trait (calving interval, CI) and 68,514 records for a survival trait (direct survival, referred to as survival for simplicity). The phenotypes were corrected for the fixed effects which were herd-year-season, parity and age for survival, and herd-year-season, parity, age at first calving and month of calving for CI, as well as the random permanent environment effect [18-20]. As some cows had multiple records (i.e., multiple survival scores or CIs), the corrected phenotypes were averaged per cow and weighted based on number of records according to [21] in our models. These cows had the same SNP genotype data as described above in our simulation study. Model performance was evaluated using five-fold cross-validation for both traits. We chose fertility and survival because they are known to be related to fitness and are affected by dominance effects from conserved regions [22].

### Genetic mixed model with dominance effects (BayesR3AD)

The genetic mixed effects model was applied to estimate additive and dominance genetic variances. Both additive and dominance genotype matrices were constructed from the genome-wide SNP data. The animal model was

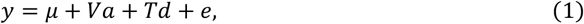

where y is an *n*_*R*_ × 1 column vector of pre-corrected phenotypic values, *n*_*R*_ is the number of phenotype records; *μ* is the overall mean; *V* is a coded additive (*n*_*R*_ × *n*_*M*_) matrix, described below to estimate additive effects across n_M_ markers, and T is a coded dominance (*n*_*R*_ × *n*_*M*_) matrix; a a vector of additive genetic effects, and d a vector of dominance effects. *e* is an *n*_*R*_ × 1 vector of residuals, where 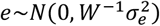 and W a diagonal weight matrix as described below.

The construction of the coded additive (*V*) and dominance (*T*) matrices follow standard quantitative genetic parameterisations. For each biallelic marker, the observed genotype for individual *i* at marker *j* was coded as *g*_*ij*_ ∈ {0,1,2}, corresponding to the reference homozygote, heterozygote, and alternative homozygote, respectively. These raw genotype counts were subsequently transformed into additive (*v*_*ij*_) and dominance (*t*_*ij*_) covariates and standardised to have mean zero and unit variance. This transformation produces centred and scaled predictors that are approximately orthogonal under Hardy-Weinberg equilibrium, improving numerical stability and enabling a clear separation of additive and dominance variance components.

Specifically, let *p*_*j*_ denote the frequency of the alternative allele at marker *j*, with *q*_*j*_ = 1 − *p*_*j*_. For genotype *g*_*ij*_ ∈ {0,1,2}, the additive covariate *v*_*ij*_ and dominance covariate *t*_*ij*_ were defined as

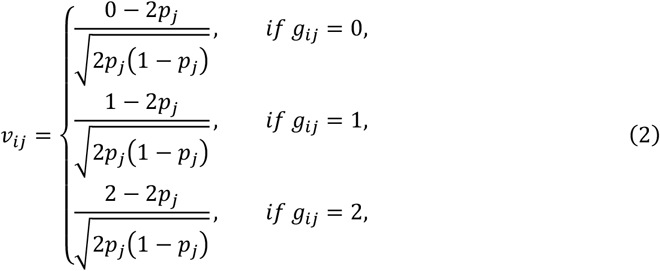

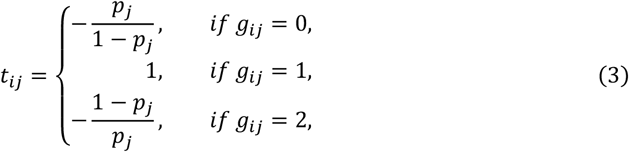

#### Mixture Proportions (π) and Variance Allocation (γ)

BayesR3AD models SNP effects using a finite mixture of normal distributions with different variances, for both additive and dominance components. For each SNP *j*, we introduce allocation indicator *k* ∈ {1, …, *K*} for both additive and dominance effects. Here *k* = 1 indicates that the corresponding SNP effect is exactly zero, while *k* ≥ 2 assign the SNP to non-zero additive and dominance variance. For each SNP *j* the additive effect (*a*_*j*_) and dominance effect (*d*_*j*_) are written as:

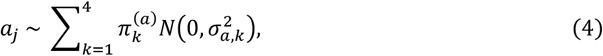

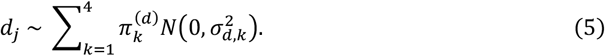

Where 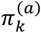 and 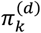 are the mixture proportions for the additive and dominance components in distribution *k*. 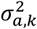 and 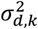 are the component-specific variances for additive and dominance effects in distribution k.

For each SNP *j*, BayesR3AD uses separate latent mixture indicators 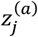 and 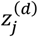 for the additive effect *a*_*j*_ and dominance effect *d*_*j*_, respectively. The indicators have separate mixture probabilities 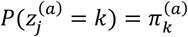 and 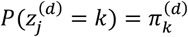, with k = 1 denoting the null component. Conditional on these indicators, a_j_ and d_j_ follow mixture-normal priors with their own variance scaling, so a SNP may have an additive effect without a dominance effect (and vice versa), and the two effects may belong to different mixture components.

By construction, the mixture proportions are probabilities and must satisfy the usual simplex constraints: 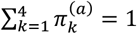 and 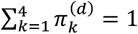, with each *π*_*k*_ ≥ 0. In BayesR3, these proportions are treated as random and updated during Markov Chain Monte Carlo (MCMC) process using a Dirichlet prior, typically Dirichlet (1, 1, 1, 1) when four mixture proportions are used [6, 9], which enforces the sum-to-one constraint and allows the model to adaptively estimate how many SNPs belong to each variance distribution.

Each mixture component is defined through a fixed variance allocation scalar *γ*_k_ that scales the genetic variance assigned to that distribution. For additive effects, we write: 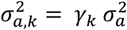, for dominance effects: 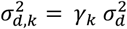 and in the standard BayesR/BayesR3 implementation, these multipliers are typically: *γ* = (0, 10^−4^, 10^−3^, 10^−2^); corresponding to a spike at zero and three non-zero variance distributions of increasing magnitude [6, 9]. An important normalisation condition links the mixture proportions and variance scaling. When genotype columns are standardised (the mean is 0 and the variance is 1), the expected per-SNP variance under the mixture prior is:

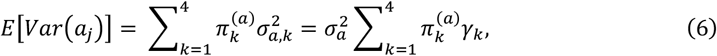

and similarly for dominance:

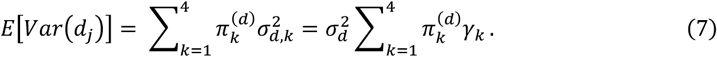

To ensure that the mixture prior is consistent with the specified genetic variances, the *γ* and *π* must be chosen (or initialised) so that the weighted sum of variance multipliers is approximately 1:

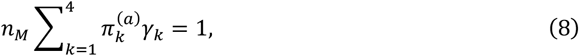

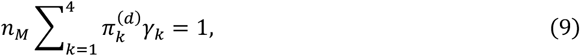

where *n*_*M*_ is the number of SNPs.

In the BayesR3 analysis, we use the same fixed γ-values for the variance distributions but treat the *π*-vectors as unknown and assign them Dirichlet priors. During MCMC, the *π*-values are updated based on the data, allowing the model to learn the proportion of SNPs in each effect-size distribution [6, 9]. The constraints ∑_*k*_ *π*_*k*_ = 1 and the fixed *γ*-values together govern how much of the total genetic variance is attributed to null vs small vs moderate vs large-effect distributions, separately for additive and dominance components.

#### Mixture weight and model entry probability

The probability that SNP *j* belongs to distribution *k* (the “in-model” vs “out-of-model” decision) is proportional to

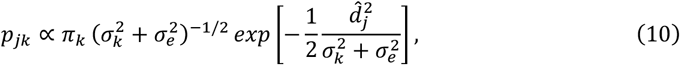

where 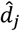 is conditional least-squares estimate of dominance effect *d*_*j*_.

The mixture weights for the non-null distributions compare each variance component to the null through the combined variance term 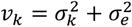 and the conditional least-squares estimate of the SNP effect. This yields the usual BayesR3 behaviour in which SNPs with very small, estimated effects tend to be allocated to the null distribution, while SNPs with larger estimated effects have higher posterior probability of belonging to small, moderate or large-effect distributions (k > 1).

#### Sampling for BayesR3 with Dominance

In this section we outline the blocked Gibbs sampling updates for the BayesR3 as specified in [9], extended to include dominance effects. Each MCMC iteration samples fixed effects *u*, additive effects *a*, dominance effects *d*, variance components, and mixture distribution allocations. In this paper, the number of MCMC iterations was 20,000 for both BayesR3 and BayesR3AD analysis. Likewise, the burning length was 10,000. Blocked Gibbs updates ensure efficient mixing, and dominance is sampled conditional on additive effects and residuals. For each iteration, the sampler updates: (i) fixed effects and residual variance, (ii) marker allocations and effects for additive and dominance components, and (iii) mixture proportions and global variance parameters. Below we describe the dominance part of sampling conditional on the remaining parameters and the data.

#### Blocked Gibbs updates for dominance

We follow the BayesR3 blocked Gibbs sampler and extend it to a model that includes both additive and dominance SNP effects. Additive allocations and effects were sampled using the same methods as [9]. For the dominance part of the model, let *T* be the dominance coded matrix containing all dominance marker codes and let *d* be the corresponding vector of dominance marker effects. The dominance contribution in block form can be written as

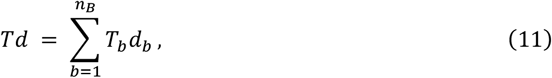

where *T*_*b*_ is the dominance submatrix for block *b* and *d*_*b*_ is the vector of dominance effects for the SNPs in that block. As in BayesR3, the dominance SNPs are partitioned into *n*_*B*_ non-overlapping blocks of size *n* (last block possibly smaller). In each outer iteration we process all dominance blocks, and within each block we perform *n* inner Gibbs updates of the block effects. With block size *n* and *m* outer iterations, each dominance SNP effect has a Markov chain of length *n*_*L*_ = *m* × *n*.

Let *ℓ* = 1, …, *m* index the outer iterations. At outer iteration *ℓ*, the dominance blocks *b* = 1, …, *n*_*B*_ are processed sequentially. On entry to dominance block *b*, we form the block right-hand side and the block cross-product matrix

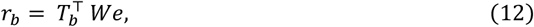

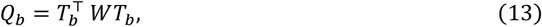

where *W* is the diagonal weight matrix and *e* is the current residual vector from the full model and *T*^⊤^ represents the transpose of *T*.

Within block *b*, we perform *n* inner iterations *i* = 1, …, *n*. At each inner iteration we loop over the dominance SNP effects in that block and update them one at a time.

Consider dominance effect *d*_*bc*_ corresponding to column *c* of *T*_*b*_. Let

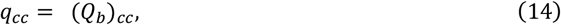

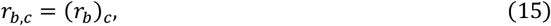

where *q*_*cc*_ represents the *c*th diagonal element of matrix *Q*_*b*_, *r*_*b,c*_ is the *c*th element of *r*_*b*_.

And let 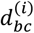 be the current value of this dominance effect at inner iteration *i*. We use the conditional least-squares estimate of the current dominance effect, conditional on all other effects,

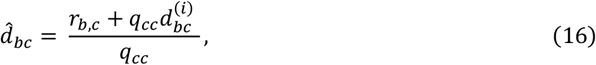

which plays the same role for dominance as the BayesR3 block wise *â*_*j*_ does for additive effects.

Dominance effects follow a finite normal mixture prior with distribution-specific variances 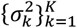 and mixture proportions 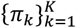. For component *k*, define

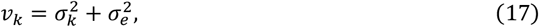

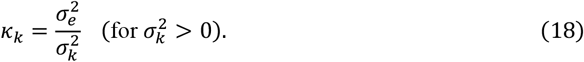

For this dominance effect *d*_*bc*_ we first compute the unnormalized weights

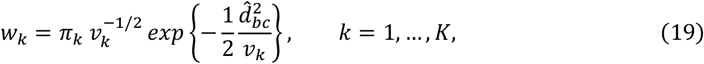

then normalise them to obtain the posterior distribution probabilities

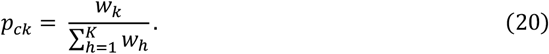

The variance distribution index *k* for this dominance effect is then sampled from a categorical distribution:

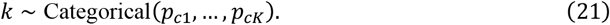

Given the sampled distribution *k*, the full conditional for *d*_*bc*_ is a univariate normal that uses the diagonal of *Q*_*b*_:

- if *k* is the null distribution (with 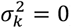), we set

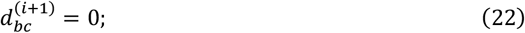
- otherwise, for 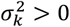,

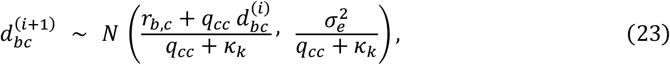

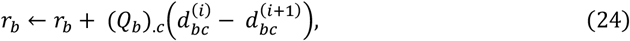

where (*Q*_*b*_)_.*c*_ is *c*th column of 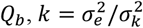 for a BayesR3 solution. The notation *x* ← *x* + 3 signifies that variable *x* is replaced by the value of *x* + 3.

We overwrite 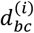 with the new draw 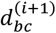, move to the next dominance effect in block *b*. Repeating this single effect update over all dominance SNPs in the block for the *n* inner iterations yield an updated dominance block vector 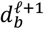 at the end of the inner cycles.

On exit from dominance block *b*, the residuals are updated in a single operation using the change in the block dominance effects:

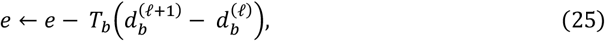

where 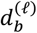 is the dominance block vector at the start of outer iteration *ℓ*. This matches the BayesR3 strategy of updating residuals once per block, while here it is applied specifically to the dominance matrix *T* and effects *d*.

#### Updating variance components of dominance effects

After each Gibbs iteration, the variances of dominance effects were sampled using inverse scaled Chi-square distributions via

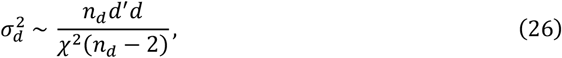

where *χ*^2^(*x*) is a Chi-square distribution with x degrees of freedom, *n*_*d*_ is the number of SNPs in dominance distribution in the model for this iteration which equals to 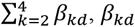 is the number of SNPs in the *k*th distribution in dominance distribution.

#### Updating dominance mixture variances and proportions

After all dominance blocks have been processed once, one outer iteration is complete. At the end of outer iteration *ℓ*, and after the global dominance SNP variance 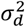 has been sampled, the distribution-specific dominance variances are reset using the fixed scaling factors *γ*_*k*_,

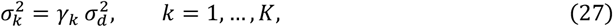

for use in the next outer iteration. The dominance mixture proportions *π* = (*π*_1_, …, *π*_*K*_) are then sampled from their Dirichlet full conditional,

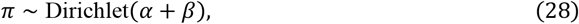

where *α* = (*α*_1_, …, *α*_*K*_) is the Dirichlet prior parameter (e.g., *α*_*k*_ = 1), and *β* = (*β*_1_, …, *β*_*K*_) contains the number of dominance SNPs assigned to each distribution *k* at the end of the outer iteration. In particular, *β*_*k*_ counts the SNPs with current distribution index *k*. Dominance SNPs with *k* > 1 at a given iteration are regarded as being “in the model”, whereas those with *k* = 1 (null distribution) are regarded as being out of the model.

### Simulation model

We built a Bayesian simulation program to simulate the phenotypes.

Let *y* be the *n* × 1 vector of phenotypes measured on *n* individuals. We consider a linear model with marker-derived additive and dominance random effects:

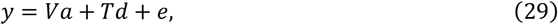

where *y* is an *n*_*R*_ × 1 column vector of phenotype values, *n*_*R*_ is the number of records; *V* is a coded genotype (*n*_*R*_ × *n*_*M*_) matrix, as constructed in the methods, representing the estimated genotypes of each individual across *n*_*M*_ markers, and *T* is a coded dominance (*n*_*R*_ × *n*_*M*_) matrix; *a* was the vector of additive genetic effects, and *d* was the vector of dominance effects; *e* is the *n* × 1 vector of residuals.

Consider one locus (*j*th SNP) for individual *i*, additive values *v*_*ij*_ and dominance value *t*_*ij*_ are coded as the same as the genetic mixed model with dominance effects, which *v*_*ij*_ and *t*_*ij*_ are scaled and centred such that each locus has a mean of 0 and standard deviation of 1.

Then we sampled the additive and dominance effects at each SNPs via:

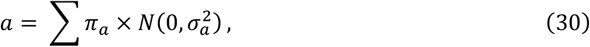

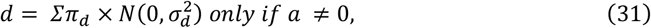

We set the constraints that only when the additive effects are non-zero, dominance effects exist.

Then we constructed genetic value *g* for each animal via

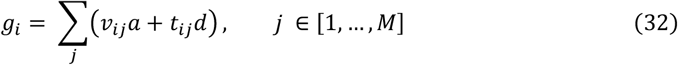

where *v*_*ij*_ is the additive genetic value of animal *i* at SNP *j* and *t*_*ij*_ is the additive genetic value of animal *i* at SNP *j, i* ∈ [1, …, *N*], *N* is the number of animals, and *j* ∈ [1, …, *M*], *M* is the number of SNPs.

After this, we calculated the genetic variance 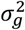 and then calculated the environmental variance via

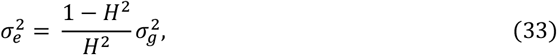

where *H*^2^ is the broad-sense heritability. Then we calculated the environmental effects via 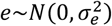. The simulated phenotypes *y* are calculated from *y* = *g* + *e* for each animal.

Using the simulation approach described above, we simulated phenotypes for 227,942 Holstein cows using their real genotypes (74,626 SNPs). We simulated two versions of phenotypes: 1) including only additive effects with additive variance 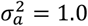, and 2) including both additive and dominance effects with additive variance 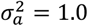 and dominance variance 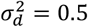. The broad-sense heritability of traits simulated was set to 0.44. Each of these phenotypes was evaluated using BayesR3, which only analyses for additive effects, and BayesR3AD, which analyses the data for both additive and dominance effects, to assess both predictive performance and the ability of BayesR3AD to recover true genetic variance components.

To ensure that the simulated genetic architecture reproduced the target additive and dominance variances, we explicitly specified the mixture proportions for the SNP effects. For additive effects, the mixture proportions were set to 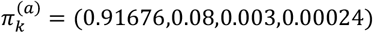. For dominance effects, proportions were specified subject to dominance-specific constraints as 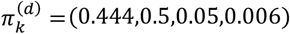, corresponding to overall SNP proportions of approximately (0.95415, 0.0416, 0.0042, 0.00005). Component variances were scaled using fixed variance allocation scalar *γ*_*k*_ = (0, 10^−4^, 10^−3^, 10^−2^). Additive and dominance genotype covariates were encoded following the breeding parameterisation of Vitezica et al. [13] and subsequently standardised to mean zero and unit variance. Under this specification, the SNP-level mixture distributions collectively reproduced the desired additive and dominance genetic variance components. When we run the BayesR3 and BayesR3AD models using the simulated phenotypes, the same additive and dominance mixture proportions as the simulation are set as the initial mixture proportion values for the first iteration in BayesR3.

### Inbreeding depression (ID)

To assess the contribution of dominance effects and to validate their biological relevance, inbreeding depression (ID) was quantified for Holstein calving interval and survival. Inbreeding depression was defined as the expected change in phenotype resulting from increased homozygosity and was computed as

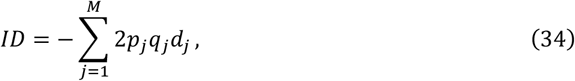

where *j* indexes SNPs (*j* = 1, …, *M*), *p*_*j*_ and *q*_*j*_ = 1 − *p*_*j*_ denote the frequencies of the reference (A) and alternative (a) alleles at locus *j*, respectively, and *d*_*j*_ is the dominance effect at that locus. Note that this unscaled dominance effect (*d*_*j*_) is different from that used in our simulations and analysis.

Under the BayesR3AD model, genotypic values were parameterised using allele-frequency–scaled additive and dominance effects as

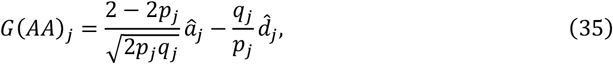

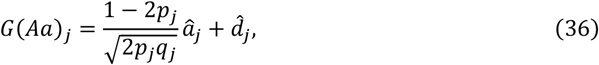

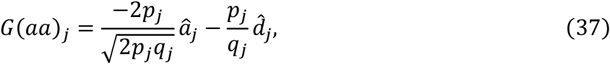

where *â*_*j*_ and 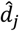 are the posterior mean estimates of the additive and dominance effects, respectively.

In classical quantitative genetics, the dominance effect *d*_*j*_ is the deviation of the heterozygote from the mid-point of the two homozygotes,

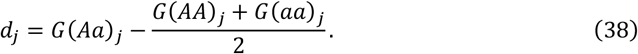

Substituting the above parameterisation yields

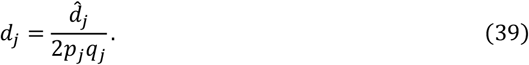

Consequently, the inbreeding depression simplifies to

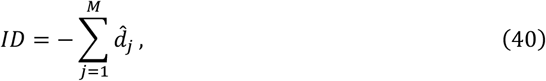

allowing ID to be computed directly from the estimated dominance effects.

### Prediction accuracy and bias

To evaluate predictive performance for both Holstein calving interval and survival, we calculated prediction accuracy and regression slope using two standard validation metrics. Prediction accuracy was defined as the Pearson correlation between the predicted genetic values and the phenotypes,

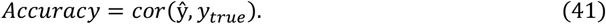

To assess potential inflation or deflation of predictions, we also estimated the bias of the true phenotype on the predicted value,

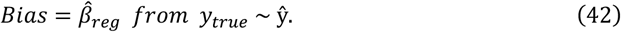

An unbiased predictor should yield a value close to one, while bias below or above one indicates over-shrinkage or under-shrinkage, respectively.

## Results

### Analysis of simulated data

#### Variance and mixture proportion results

The simulation results demonstrate the clear importance of modelling dominance when dominance contributes to phenotypic variation. In the additive-only simulation scenario (Table 1), both BayesR3 and BayesR3AD produced estimates close to the expected additive variance. Although this dataset contained no dominance effects, the inclusion of dominance in the model (BayesR3AD) only estimated minimal dominance effects and is reflected by 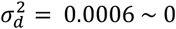.

**Table 1.**
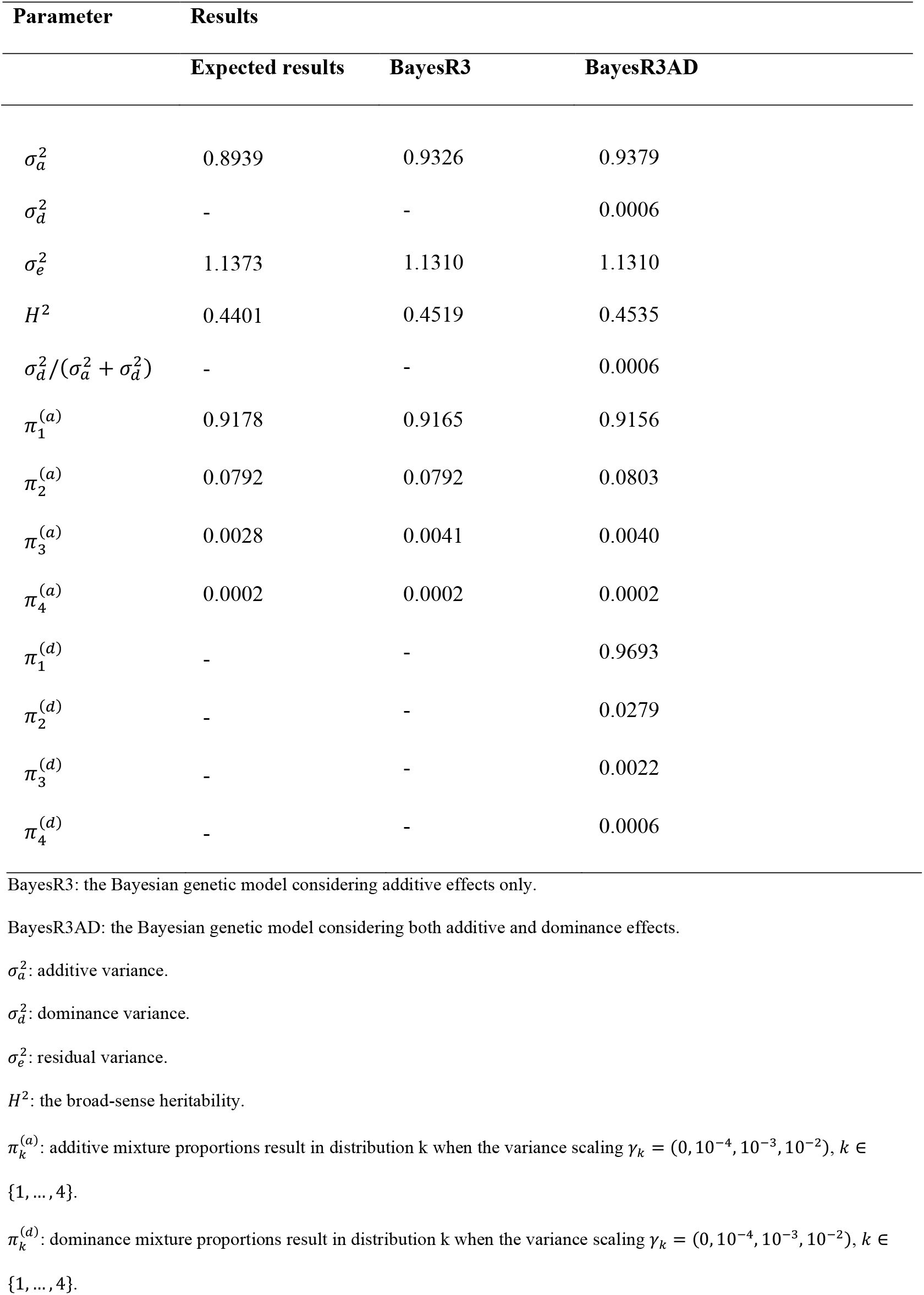
Variance and mixture proportion from BayesR3 and BayesR3AD using simulated phenotypes with additive effects.

For the additive mixture proportion results (Table 1), both BayesR3 and BayesR3AD recovered the expected additive mixture proportions with high accuracy across the first, second, and fourth mixture distributions. The only minor deviation was observed in the third distribution, where both models slightly overestimated the proportion relative to the expected value (0.004 vs. 0.0028). As anticipated, the dominance mixture proportions in BayesR3AD were extremely small across all dominance classes (0.9693, 0.0279, 0.0022, 0.0006). These values reflect the absence of dominance effects in the simulated data and demonstrate the model’s conservative behaviour when dominance is not present.

When phenotypes were simulated with both additive and dominance effects (Table 2), the advantages of explicitly modelling dominance became substantial. BayesR3AD accurately recovered the dominance variance 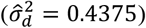 in close agreement with the true simulated value (0.4359), and it correctly partitioned additive and environmental variance components. In contrast, the additive-only model (BayesR3) was forced to absorb dominance effects into the residual, yielding a markedly inflated estimate of environmental variance (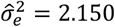 compared with the true value of 1.745) and decreasing heritability by 12% (*H*^2^ = 0.4380 → 0.3164). Notably, the inflated residual variance approximates the sum of the true environmental and dominance variances, illustrating the expected confounding that arises when dominance is omitted. Such misspecification also biases heritability estimates downward, as the additive component no longer reflects the true genetic contribution.

**Table 2.** Variance and mixture proportion from BayesR3 and BayesR3AD using simulated phenotypes with additive and dominance effects.

| Parameter | Results |  |  |
| --- | --- | --- | --- |
|  | Expected results | BayesR3 | BayesR3AD |
| $\sigma_a^2$ | 0.9242 | 0.9952 | 0.9598 |
| $\sigma_d^2$ | 0.4359 | - | 0.4375 |
| $\sigma_e^2$ | 1.7456 | 2.1500 | 1.7350 |
| $H^2$ | 0.4380 | 0.3164 | 0.4461 |
| $\sigma_d^2/(\sigma_a^2 + \sigma_d^2)$ | 0.3205 | - | 0.3131 |
| $\pi_1^{(a)}$ | 0.9178 | 0.9110 | 0.9150 |
| $\pi_2^{(a)}$ | 0.0792 | 0.0851 | 0.0807 |
| $\pi_3^{(a)}$ | 0.0028 | 0.0037 | 0.0041 |
| $\pi_4^{(a)}$ | 0.0002 | 0.0002 | 0.0002 |
| $\pi_1^{(d)}$ | 0.9558 | - | 0.9520 |
| $\pi_2^{(d)}$ | 0.0396 | - | 0.0435 |
| $\pi_3^{(d)}$ | 0.0042 | - | 0.0041 |
| $\pi_4^{(d)}$ | 0.0005 | - | 0.0004 |

BayesR3: the Bayesian genetic model considering additive effects.
BayesR3AD: the Bayesian genetic model considering both additive and dominance.
$\sigma_a^2$ : additive variance.
$\sigma_d^2$ : dominance variance.
$\sigma_e^2$ : residual variance.
$H^2$ : the broad-sense heritability.

The starting additive mixture proportions for both BayesR3 and BayesR3AD were set to (0.91676, 0.08, 0.003, 0.00024), while the starting dominance mixture proportions were specified as (0.95655, 0.039, 0.004, 0.00045). As shown in Table 2, the final additive mixture proportions inferred by both models closely matched the expected values. Likewise, the dominance mixture proportions estimated by BayesR3AD accurately recovered the expected distribution across all four mixture components. To assess sensitivity to the mixture proportion specification, we repeated the analysis using alternative starting mixture proportions that differed substantially from the true values. The final estimates converged to the expected results, demonstrating the robustness of BayesR3AD and its ability to recover heterogeneous dominance effect-size distributions.

#### Estimated SNP effects

In Figure 1 Manhattan plots present the expected and estimated SNP effects when the underlying genetic architecture comprises purely additive effects. Panel 1a shows the true additive effects simulated across chromosomes, exhibiting sparse but correctly located non-zero signals. Figure 1b displays the additive effects estimated using BayesR3 (additive-only model). The pattern closely reproduces the simulated architecture, with major peaks correctly recovered and background noise remaining small. The correlation between the true and estimated additive effects using BayesR3 was 0.7615. This demonstrates that BayesR3 performs well when the model specification matches the true architecture.

**Figure 1.**
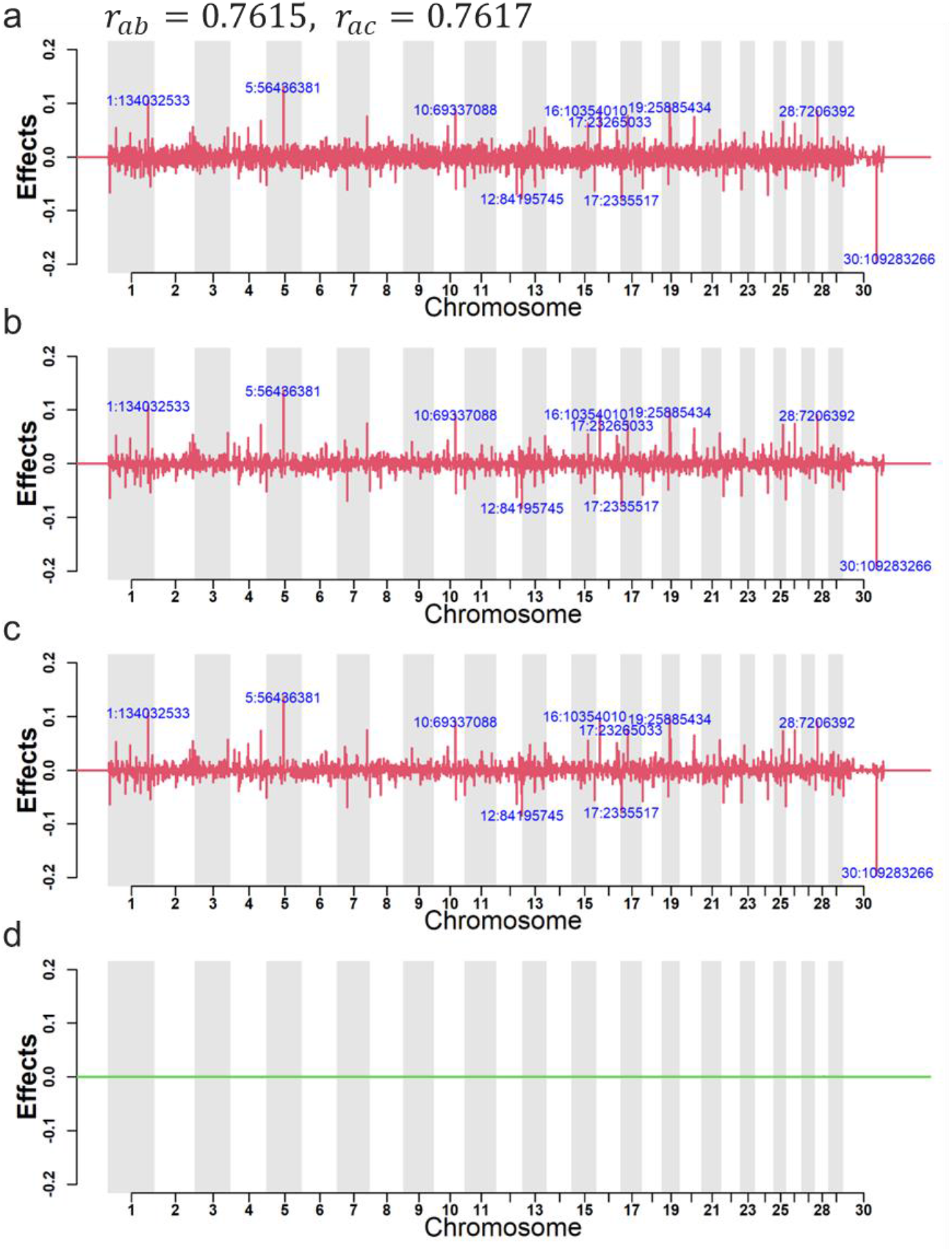
Manhattan plot comparison when simulation is additive-only. a: expected additive effects. b: estimated additive effects from BayesR3. c: estimated additive effects from BayesR3AD. d: estimated dominance effects from BayesR3AD. Top 10 SNPs according to effect magnitude are highlighted in blue text. The r values above the panel a are Pearson’s correlation values. r_ab_ is the correlation between true and estimated additive effects from BayesR3, r_ac_ is the correlation between true and estimated additive effects from BayesR3AD.

Figure 1c shows the additive effects estimated using BayesR3AD, which additionally includes dominance mixture components. The additive signals remain nearly identical to those obtained from BayesR3 with a correlation of 0.7617, indicating that incorporating dominance terms does not distort additive-effect inference when dominance is absent. Importantly, the dominance panel (Figure 1d) shows only noise-level estimates around zero with no structure or large-effect outliers. This confirms that BayesR3AD appropriately shrinks dominance effects to zero when they are not supported by the data, demonstrating the model’s desirable self-regularising behaviour.

**Figure 2** shows the performance of BayesR3AD when both additive and dominance effects are present in the simulated architecture. Panels 2a and 2b display the true additive and dominance effects, respectively. Both components include moderate, locus-specific peaks representing causal variants.

**Figure 2.**
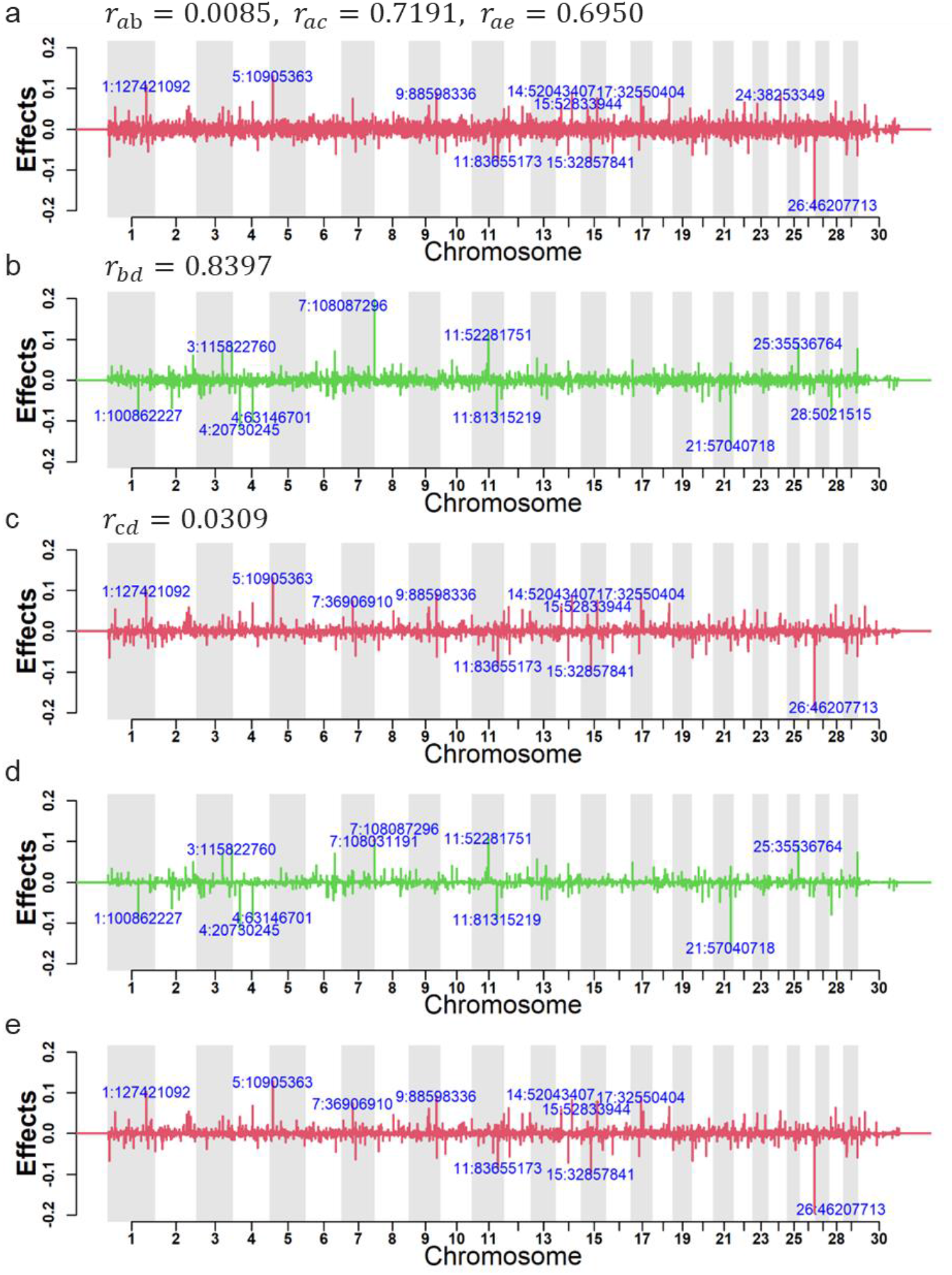
Manhattan plot comparison when simulation contains additive and dominance. a: expected additive effects; b: expected dominance effects; c: estimated additive effects from BayesR3AD; d: estimated dominance effects from BayesR3AD; e: estimated additive effects from BayesR3. Top 10 SNPs according to effect magnitude are highlighted in blue text. The r values above the panel a and b are Pearson’s correlation values. r_ac_ is the correlation between true and estimated additive effects from BayesR3AD, r_ae_ is the correlation between true and estimated additive effects from BayesR3, r_bd_ is the correlation between true and estimated dominance effects from BayesR3AD.

Panels 2c and 2d show the additive and dominance effects estimated by BayesR3AD. The model successfully recovers the major additive peaks, with patterns closely resembling those in the expected additive effects (2a) that the correlation of additive effects was 0.7191 and the correlation of dominance effects was 0.8397 between the true and estimated values from BayesR3AD. More importantly, the dominance panel (2d) captures the true dominance-effect signals with clear peaks at the correct chromosomal positions, illustrating accurate separation of additive and dominance contributions. This demonstrates that the mixture prior in BayesR3AD effectively identifies and shrinks effects into the correct additive or dominance components when both types of genetic signal are present.

Among the top 10 SNPs ranked by effect size, nine were shared between the expected and estimated additive effects (Panels 2a and 2c), and likewise nine were shared between the expected and estimated dominance effects (Panels 2b and 2d). This high concordance between simulated and posterior top loci indicates that the model accurately recovers major effect signals and supports reliable fine-mapping performance.

Panel 2e displays the additive effects estimated by the BayesR3 (additive-only model), which recover the same major additive peaks observed in the expected additive effects. The correlation between the true and estimated additive effects under BayesR3 was 0.695, indicating a strong but reduced concordance relative to BayesR3AD.

The Manhattan plot results using simulated data demonstrate that BayesR3AD behaves conservatively in the absence of dominance and does not introduce spurious dominance effects, while preserving accurate estimation of additive architecture. BayesR3AD accurately decomposes additive and dominance genetic variance when both components are present, while the BayesR3 model confounds the two sources of signal. This supports the use of BayesR3AD for traits where non-additive effects are expected to contribute meaningfully to genetic architecture.

#### Prediction accuracy and bias

**Table 3** shows the prediction accuracy and bias results using 68,383 validation individuals. When only additive effects were simulated, both BayesR3 and BayesR3AD achieved identical prediction accuracy (0.6306), indicating no penalty for fitting dominance. When dominance effects were simulated, BayesR3AD achieved an increased accuracy of 0.6144 vs. 0.5133 for BayesR3 (absolute +0.1011; ≈19.7% relative). The BayesR3AD-A results, computed using only the additive component from BayesR3AD, were nearly identical to BayesR3 (accuracy 0.5169 vs 0.5133; bias 1.0047 vs 0.9948), indicating that inclusion of dominance does not affect additive effect estimation.

**Table 3.**
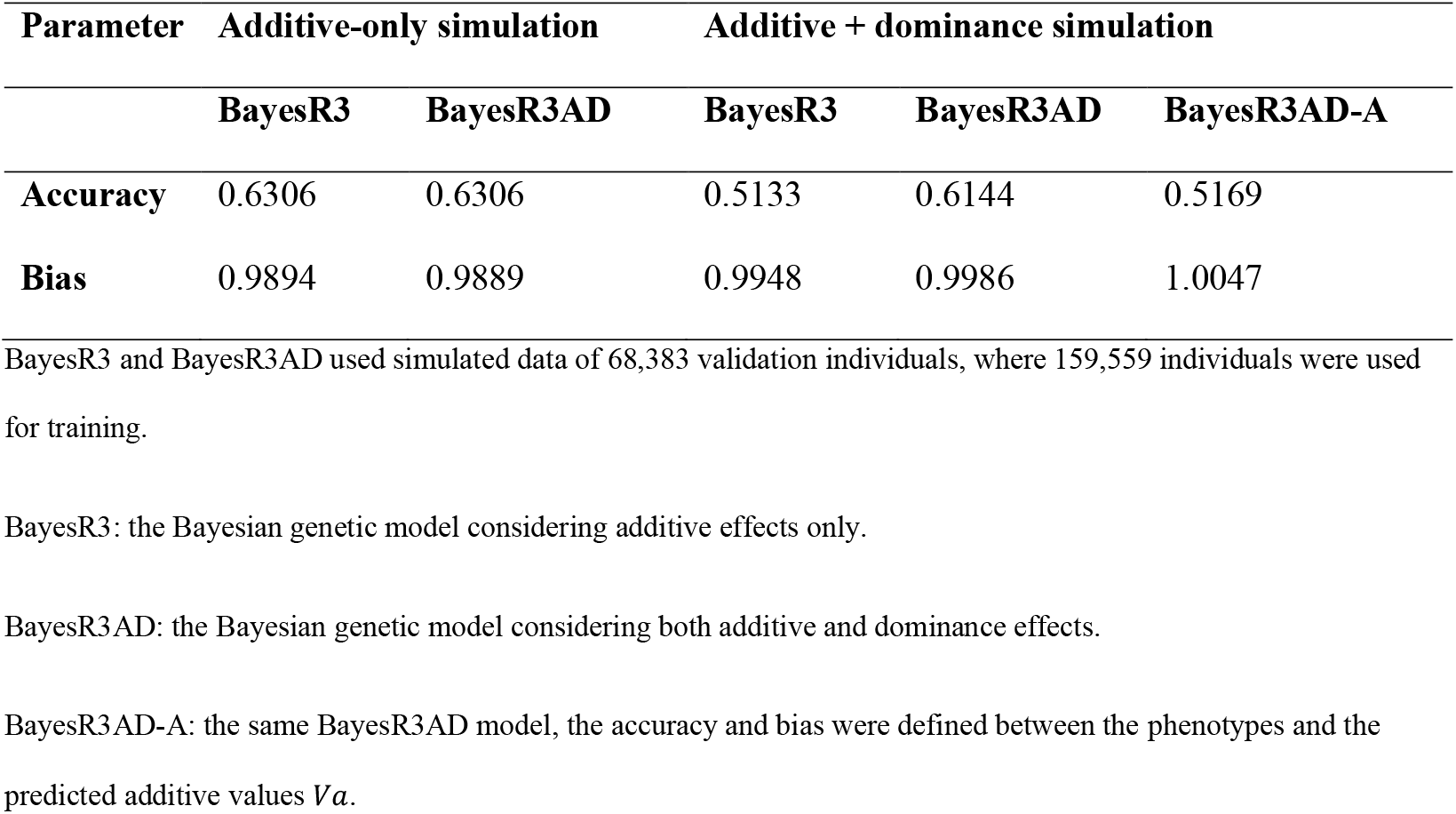
Prediction accuracy and bias.

Prediction bias shown in **Table 3** was minimal for both models across simulation scenarios, with regression slopes close to unity. In the additive-only setting, BayesR3 and BayesR3AD showed nearly identical calibration. When dominance was present, BayesR3AD achieved slightly improved calibration, supporting accurate and unbiased prediction under the extended model.

#### Computational Performance

Computational performance was evaluated for both models under the two simulation scenarios. In the additive-only simulation, BayesR3 required 15 min and 47.16 GiB of RAM, whereas BayesR3AD required 22 min and 92.82 GiB. In the additive and dominance simulation, runtimes were 18 min and 31 min for BayesR3 and BayesR3AD, respectively, with memory usage of 47.13 GiB and 92.81 GiB. Thus, incorporating dominance approximately doubled memory requirements and increased runtime by 40-70%, while remaining computationally tractable for datasets of this scale.

### Analysis of real data: Holstein Fertility (Calving Interval) and Cow Survival

#### Variance and mixture proportion results

The dominance variances estimate for calving interval and survival (**Table 4**) are small and this is consistent with the estimates for individual SNPs. For calving interval, the contribution of dominance variance to the genetic variance was 1.4%. For survival, dominance effects accounted for approximately 3% of the total genetic variance, again indicating a small contribution. Importantly, the inclusion of dominance did not destabilise estimates of additive or residual variance, and the estimated error variance 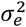 represents the residual variance for cows with a single observation.

**Table 4:**
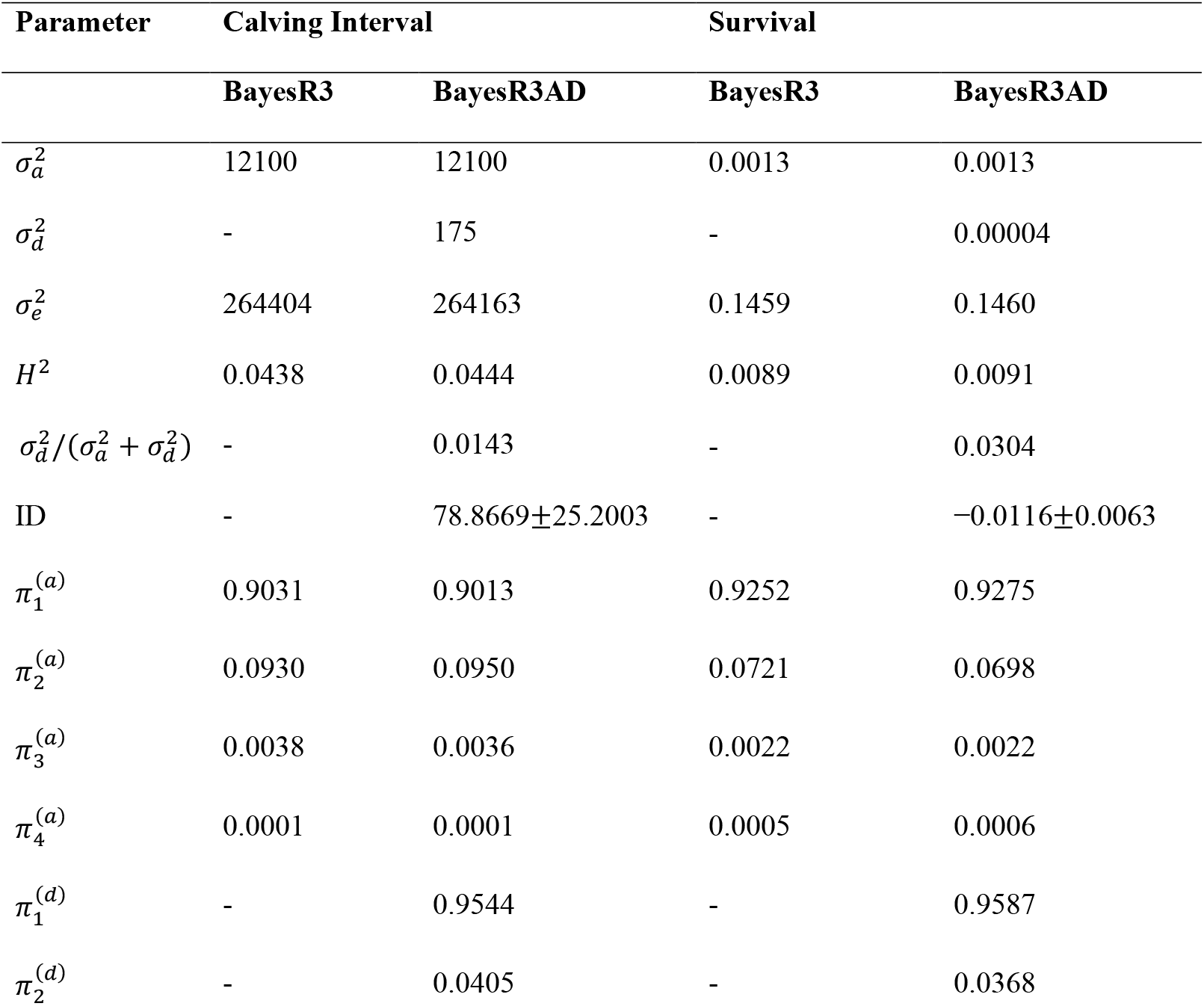
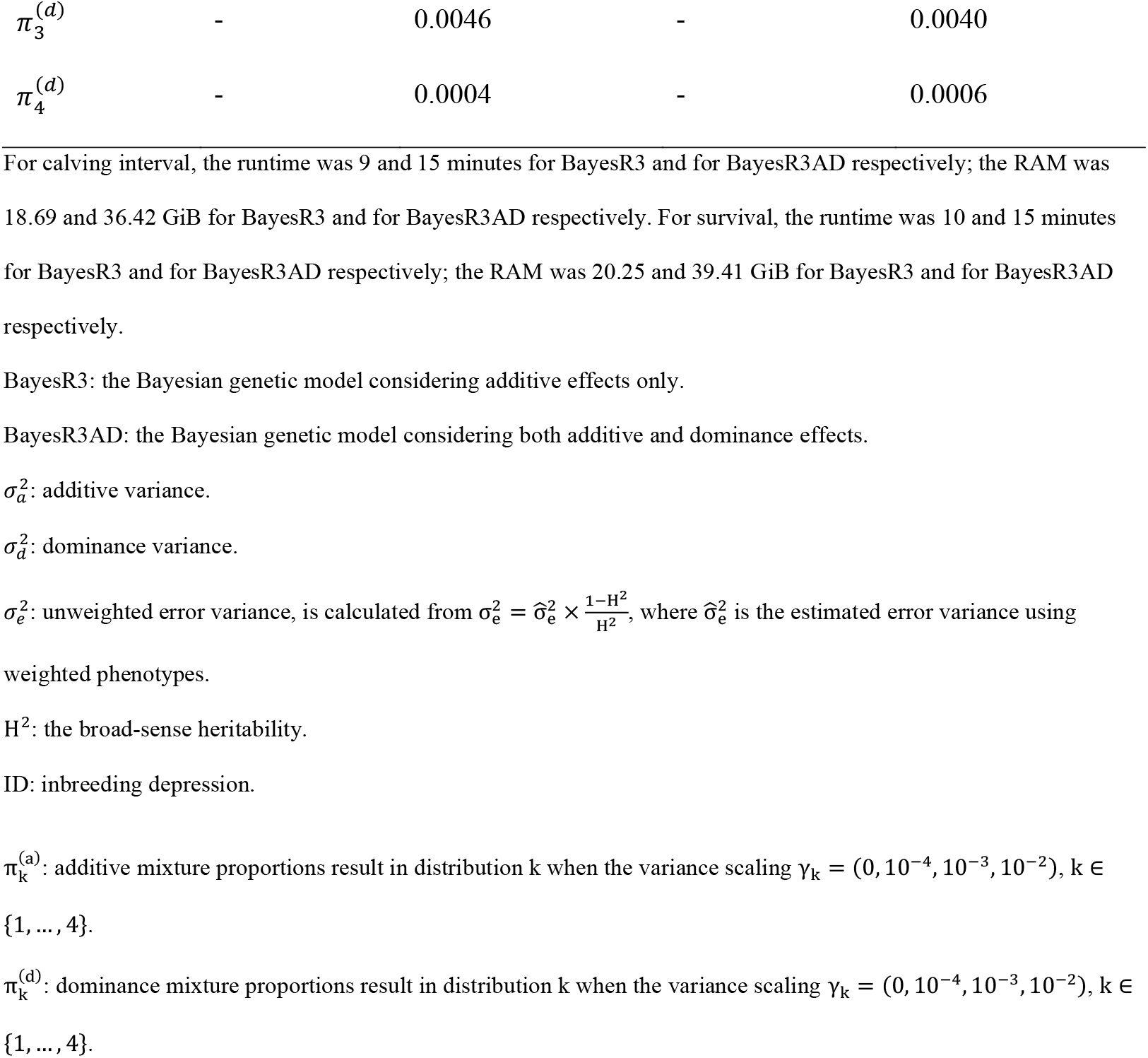
Variance and mixture proportion for calving interval and survival recorded in 63,378 and 68,514 Holstein cows respectively.

The estimated inbreeding depression for calving interval was 78.87 (SE = 25.20), whereas for survival it was −0.0116 (SE = 0.0063). The direction of these estimates is biologically consistent, indicating that increased inbreeding is associated with a longer calving interval and reduced survival. Both effects reflect a deterioration in fertility-related performance, supporting the expectation that increased homozygosity adversely affects fitness traits in dairy cattle [23]. The magnitude of the estimates is likely conservative, reflecting the shrinkage of dominance effects inherent to the Bayesian mixture prior in BayesR3AD.

**Table 4** also presents the mixture proportion estimates obtained from Holstein calving interval and survival phenotypes. The starting additive mixture proportion values in both BayesR3 and BayesR3AD were set to (0.91676, 0.08, 0.003, 0.00024) and the starting dominance mixture proportions were set to (0.95655, 0.039, 0.004, 0.00045).

The final mixture proportions for both traits remained close to the starting values, consistent with the strong sparsity typical of quantitative traits in which most loci have negligible or very small effects. To assess sensitivity to initialisation and chain length, we repeated the analysis using alternative starting mixture proportions and extended MCMC chains (200,000 iterations). Across all runs, the additive mixture proportions converged to nearly identical posterior estimates regardless of starting values, indicating good identifiability of the additive effect-size distribution. For dominance, the posterior allocations were also largely stable, with most loci assigned to the null (first) mixture component, resulting in near convergence of the dominance mixture proportions.

#### Estimated SNP effects

For calving interval, the Manhattan plots constructed from posterior inclusion probabilities (PIPs) (Figure 3) and posterior mean SNP effects (Figure 4) reveal a concentrated cluster of both additive and dominance signals on BTA18. The additive Manhattan plot (Figure 4a) highlights a major additive signal at approximately 57.82 Mb, consistent with previously reported fertility QTL in this region [24]. A second prominent additive peak occurs near 44.37 Mb. The dominance Manhattan plot (Figure 4b) reveals the same distinct peak at around 44.37 Mb on BTA18, where the estimated negative dominance effect indicates that homozygous recessive genotypes are associated with prolonged calving intervals. This signal lies close to the CHST8 gene region, previously implicated in recessive fertility biology [25].

**Figure 3.**
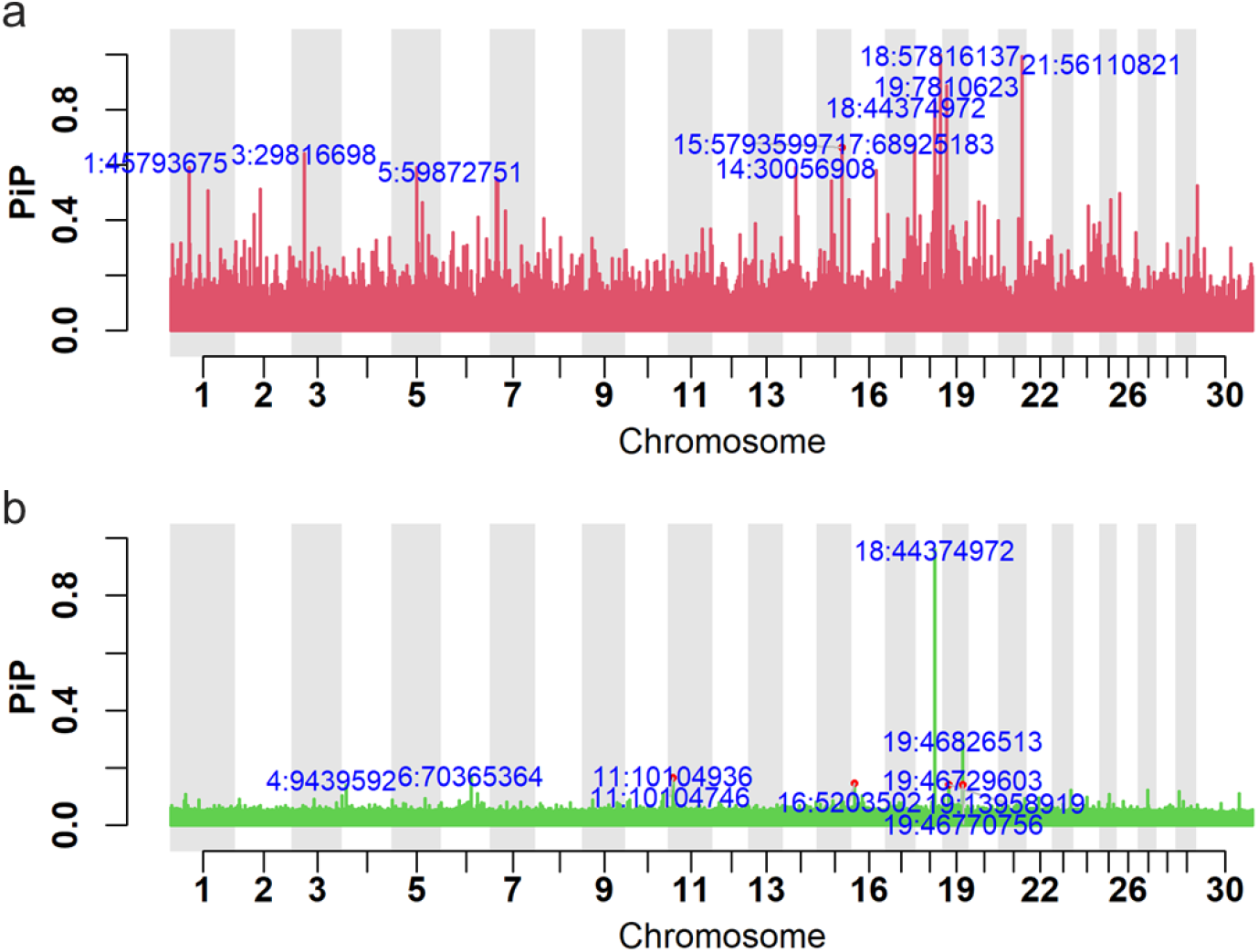
Manhattan plot of BayesR3AD posterior inclusion probabilities (PiP) for the calving interval trait. (a) shows the additive PiP and (b) the dominance PiP. In both plots the top 10 SNPs according to PiP magnitude are highlighted in blue text.

**Figure 4.**
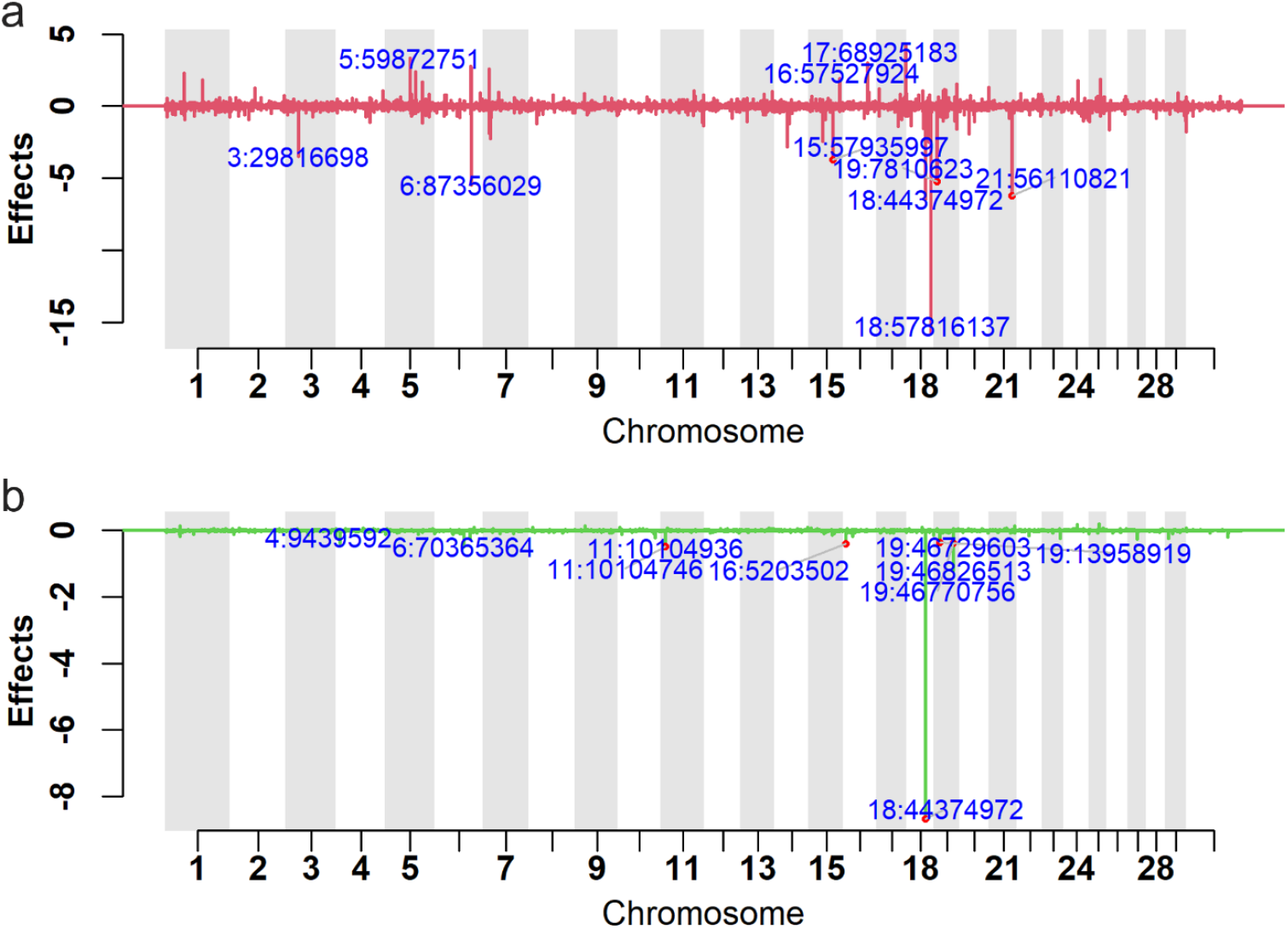
Manhattan plot of BayesR3AD estimated additive (a) and dominance (b) effects for the calving interval trait. In both plots the top 10 SNPs according to effect magnitude are highlighted in blue text.

For survival, the dominance Manhattan plot (**Figure 5**b and **Figure 6**b) similarly identifies a locus on BTA18, with a signal centered near 43.15 Mb. The estimated dominance effect on this region is positive, indicating an upward deviation of heterozygotes from the additive expectation. The region lies near the RGS9BP gene, which encodes a regulator of G-protein signaling; variants in this region have been highlighted as candidate loci in multi-trait GWAS of cattle reproductive traits, including those linked to fertility pathways (e.g., GnRH signaling and related endocrine processes) in Bos indicus populations [26, 27]. Although RGS9BP itself has not been functionally validated for survival traits, its involvement in intracellular signaling pathways and its proximity to reproduction-associated signals make it a candidate for further investigation.

**Figure 5.**
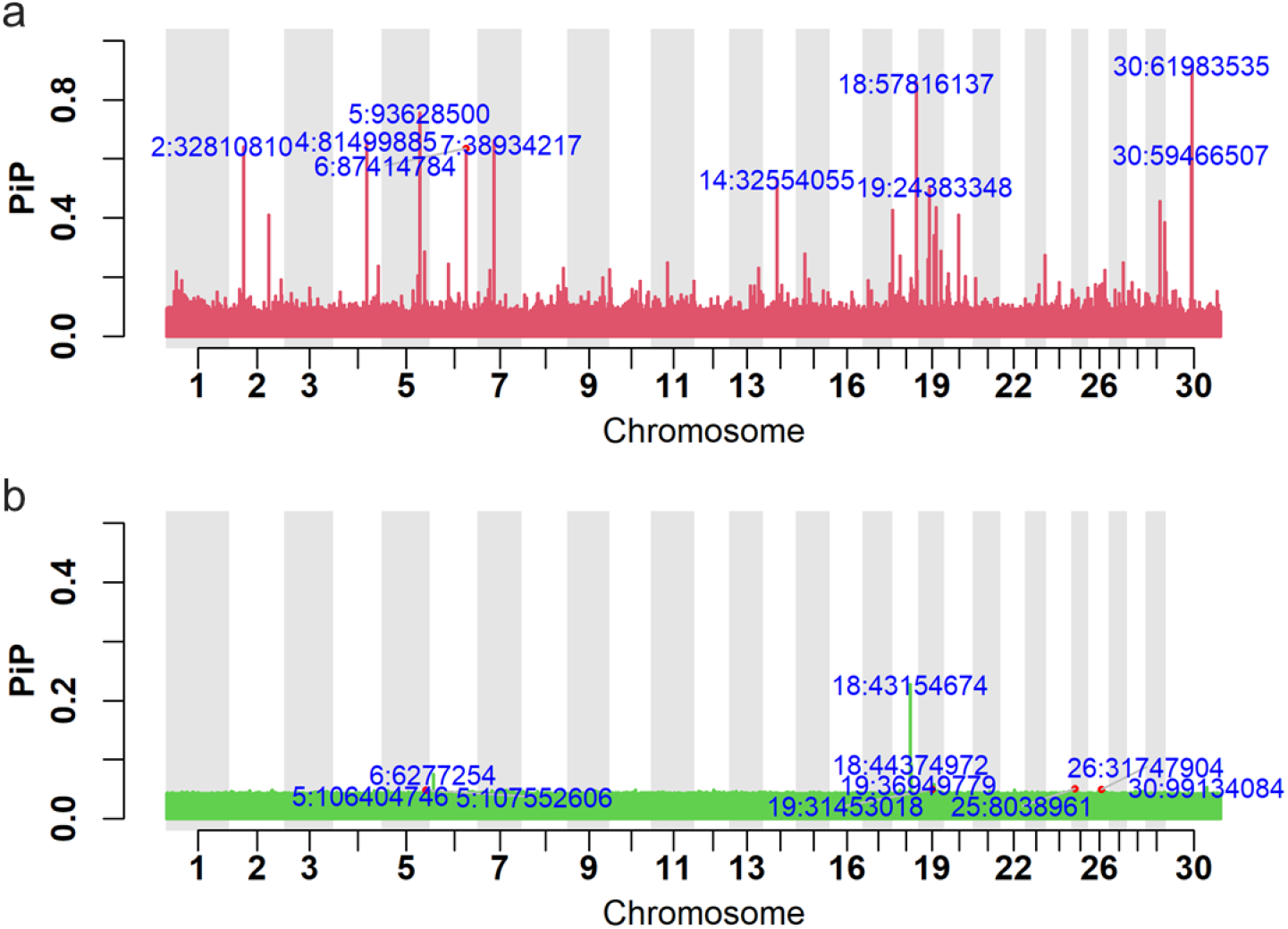
Manhattan plot of BayesR3AD posterior inclusion probabilities (PiP) for the survival trait. (a) shows the additive PiP and (b) the dominance PiP. In both plots the top 10 SNPs according to PiP magnitude are highlighted in blue text.

**Figure 6.**
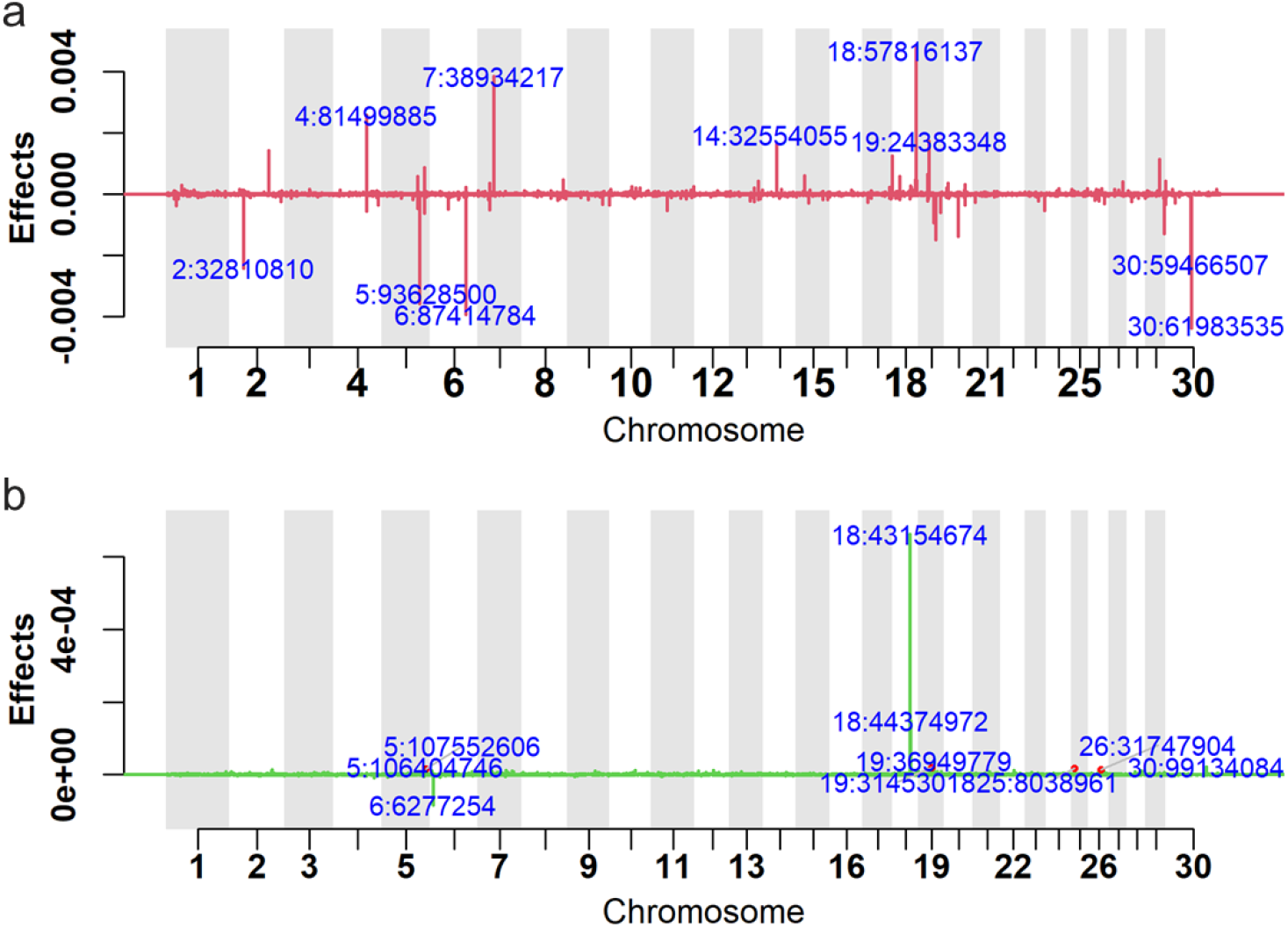
Manhattan plot of estimated additive (a) and dominance (b) effects for the survival trait using BayesR3AD. In both plots the top 10 SNPs according to effect magnitude are highlighted in blue text.

#### Prediction accuracy and bias

We used 5-fold cross validation to test the accuracy of predictions using the BayesR3 and BayesR3AD models (**Table 5**). For calving interval, using 74,626 SNPs and phenotypes from 63,378 Holstein cows, the BayesR3 model achieved a prediction accuracy of 0.243, whereas the BayesR3AD model reached 0.245. The improvement in accuracy was small, consistent with the low dominance variance estimated for this trait. A similarly small difference was observed for survival. Overall, BayesR3AD produced prediction accuracies that were comparable to BayesR3, while accommodating dominance effects without compromising additive inference.

**Table 5.** Prediction accuracy and bias using 5-fold cross validation.

| Parameter | Calving Interval |  | Survival |  |
| --- | --- | --- | --- | --- |
|  | BayesR3 | BayesR3AD | BayesR3 | BayesR3AD |
| Accuracy | 0.243 ± 0.010 | 0.245 ± 0.010 | 0.190 ± 0.007 | 0.191 ± 0.007 |
| Bias | 1.103 ± 0.060 | 1.107 ± 0.062 | 1.820 ± 0.087 | 1.819 ± 0.094 |

Prediction accuracy and bias are presented as mean ± standard deviation (SD).

## Discussion

Being able to identify and utilise dominance effects is important in both animal and plant breeding. While our new method is demonstrated using animal data, it can be directly applied to other species. A key strength of BayesR3AD is its robustness as a Bayesian mixture approach. It integrates dominance effects directly into the Bayesian mixture modelling of SNP effects, enabling joint estimation of additive and dominance variance components. The model inherits the adaptive shrinkage behaviour characteristic of BayesR3: when dominance effects are absent, the dominance variance is estimated to be near zero, preventing overfitting and preserving accurate estimation of additive heritability. Conversely, when dominance is present in the data, BayesR3AD activates the appropriate mixture components, allowing the dominance variance to be captured without biasing the additive component. This adaptive capability is central to the model’s stability across different genetic architectures.

The simulations demonstrate that correctly modelling dominance is crucial for accurate prediction when dominance contributes meaningfully to trait variation. When dominance effects were included in the phenotype simulation but omitted from the prediction model, a substantial proportion of the genetic variance was misallocated to the residual term, leading to reduced accuracy. The nearly ten‐percentage‐point improvement in prediction accuracy when applying BayesR3AD to dominance‐simulated traits illustrates its practical value for breeding programs aiming to predict total genetic merit rather than additive merit alone.

The analyses using real data for calving interval and survival suggest that dominance contributes only modestly to the overall genetic architecture of these two traits. The observed gains in predictive accuracy from BayesR3AD over BayesR3 were very small, and the regression slopes of true on predicted phenotypes remained close to those from the additive model. These patterns indicate that the dominance component is neither overfitted nor inflated, and that the extended model largely preserves the additive predictions that drive selection decisions. Previously, we observed much stronger dominance effects enriched in conserved sites for fertility and survival, compared to sites not conserved [28]. Therefore, a future direction could be to model different genomic regions in the BayesR3AD model to improve the prediction accuracy. However, the results do find inbreeding depression for both traits and locate at least one deleterious recessive mutation of moderate effect. The finding that dominance variance is near zero despite the existence of inbreeding depression is not unexpected when many variants contribute to the trait.

Beyond prediction, BayesR3AD provides rich biological interpretation. By delivering posterior estimates and inclusion probabilities for both additive and dominance effects at each SNP, the model supports locus-specific inference, fine mapping, and the identification of regions where heterozygote effects are particularly important.

The BTA18 region is a key genomic area in Holstein cows associated with several important functional traits, including udder health, calving difficulty, fertility, longevity, and body conformation [29-31]. Liang et al. [25] reported a broad additive association region on BTA18 (43.1-59.7 Mb), together with significant dominance effects in the CEBPG - PEPD - CHST8 gene region (43.8-44.18 Mb), from a large-scale GWAS of fertility traits in Holsteins. The posterior signals on BTA 18 identified by BayesR3AD overlap these intervals but more precisely resolved two apparently independent QTL, with a major additive peak at ∼57.82 Mb and a second locus at ∼44.37 Mb showing both additive and dominance effects.

The 57.82 Mb signal falls close to the well-established calving and fertility QTL region (≈57.32-57.62 Mb), where multiple studies have reported strong effects on direct calving traits, including a lead SNP near 57.59 Mb (e.g., [24, 32-34]). This concordance with prior GWAS supports the ability of BayesR3AD to recover validated loci while providing refined, locus-specific effect estimates within a multi-locus Bayesian framework.

A second signal corresponded to a prominent negative dominance peak on BTA18 (chr18: 44,374,972), located proximally to the CEBPG - PEPD - CHST8 gene region [25]. The concordance between our dominance signal and this previously identified fertility region suggests that non-additive genetic effects at approximately this locus contribute to fertility, with the estimated direction consistent with reduced calving interval and improved reproductive performance.

In addition to BTA18, Liang et al. identified a 0.53 Mb region on chromosome 6 (86.79-87.32 Mb) containing GC, NPFFR2, and SLC4A4, which harboured multiple top-ranked additive associations. Consistent with these findings, BayesR3AD detected several large additive effects within and near this interval, including a prominent peak at 87.35 Mb, reinforcing the relevance of this region. The agreement between BayesR3AD and established QTL reinforces the model’s capacity to recover true multi-locus additive architecture.

Collectively, these results demonstrate the capacity of BayesR3AD to recover GWAS-scale signals within a fully Bayesian multi-locus framework. Such locus-specific dominance effects are difficult to characterise using models that account for dominance exclusively through dense genomic relationship matrices, which distribute dominance variance genome-wide and lack fine-mapping resolution [12, 13]. In contrast, the mixture prior structure of BayesR3AD enables sparse dominance effects to be modelled flexibly, improving detection and interpretation of heterogeneous non-additive genetic signals.

From an applied breeding perspective, improved decomposition of genetic variance into additive and dominance components has several implications. First, it refines estimates of additive heritability and breeding values, ensuring that non-additive variation is not misattributed to the residual. Second, explicit modelling of dominance enhances predictions of total genetic merit, which is particularly relevant for mating allocation, crossbreeding schemes, and management of recessive defects. Third, the asymmetry in the model’s behaviour, conservative when dominance is negligible, and responsive when it is substantial, means that breeders can adopt BayesR3AD without risking deterioration of predictive performance for predominantly additive traits.

## Conclusions

BayesR3AD extends BayesR3 to jointly model additive and dominance SNP effects using a blocked Gibbs sampler in which additive and dominance blocks are processed sequentially, with residuals updated once per block via the block-level change in effects. Simulation studies demonstrate that when dominance contributes to trait variation, BayesR3AD recovers dominance variance and improves prediction accuracy relative to an additive-only BayesR3 model, whereas when dominance is weak the method behaves conservatively, leaving additive predictions essentially unchanged. Applications to Holstein fertility traits show that, even when dominance plays a modest role, BayesR3AD can detect biologically plausible dominance loci and produce small gains in predictive accuracy without compromising additive inference. More broadly, in other species, such as many plant breeding systems, where dominance plays more significant roles, BayesR3AD has the potential to deliver more substantial improvements in new loci discovery and genomic prediction. Together, these features make BayesR3AD a practical and biologically informative tool for breeding programs aiming to capture both additive and non-additive components of genetic architecture.

## Declarations

### Availability of data and materials

DataGene Limited (http://www.datagene.com.au/) manages the raw phenotype and genotype data of Australian dairy animals, and access to these data for research purposes may be granted upon request to DataGene. BayesR3AD software is available for research purposes on request.

### Authors’ contributions

M.E.G. and E.J.B conceived the study. H.Y. and E.J.B. developed the executables for BayesR3AD software and wrote the initial draft of the paper. E.J.B. and M.E.G. developed the equations. H.Y. ran the analysis. I.M.M. assisted with simulation studies. M.K. corrected the phenotypes. M.E.G. and R.X. oversaw the study and secured the funding. All authors revised and approved the manuscript.

## Acknowledgements

The Australian Research Council’s Discovery Projects (DP160101056, DP200100499, and DP230101352) supported M.E.G. This work is part of a project within the “DairyBio” programme and was funded by Agriculture Victoria (Melbourne, Australia), Dairy Australia (Melbourne, Australia) and the Gardiner Foundation (Melbourne, Australia). We thank Australian dairy farmers for data recording and DataGene staff, especially Dr Gert Nieuwhof for data access and processing.

